# CD56-mediated activation of human natural killer cells is triggered by *Aspergillus fumigatus* galactosaminogalactan

**DOI:** 10.1101/2023.08.18.553508

**Authors:** Linda Heilig, Fariha Natasha, Nora Trinks, Vishukumar Aimanianda, Sarah Sze Wah Wong, Thierry Fontaine, Ulrich Terpitz, Lea Strobel, François Le Mauff, Donald C. Sheppard, Sascha Schäuble, Oliver Kurzai, Kerstin Hünniger, Esther Weiss, Mario Vargas, P. Lynne Howell, Gianni Panagiotou, Sebastian Wurster, Hermann Einsele, Jürgen Löffler

## Abstract

Invasive aspergillosis causes significant morbidity and mortality in immunocompromised patients. Natural killer (NK) cells are pivotal for antifungal defense. Thus far, CD56 is the only known *pathogen recognition receptor* on NK cells triggering potent antifungal activity against *Aspergillus fumigatus*. However, the underlying cellular mechanisms and the fungal ligand of CD56 have remained unknown. Using purified cell wall components, biochemical treatments, and *A. fumigatus* mutants with altered cell wall composition, we herein found that CD56 interacts with the *A. fumigatus* cell wall carbohydrate galactosaminogalactan (GAG). This interaction induced NK cell activation, degranulation, and secretion of immune-enhancing chemokines and cytotoxic effectors. Supernatants from GAG-stimulated NK cells elicited antifungal activity and enhanced antifungal effector responses of polymorphonuclear cells. In conclusion, we identified *A. fumigatus* GAG as a ligand of CD56 on human primary NK cells, stimulating potent antifungal effector responses and activating other immune cells.

## Introduction

Invasive pulmonary aspergillosis (IPA), most commonly caused by the opportunistic mold pathogen *Aspergillus fumigatus*, is a devastating infection in immunocompromised patients. Individuals that are highly susceptible to IPA often have neutropenia and/or dysfunctional T-cell responses ^1^. This includes patients suffering from hematological malignancies, undergoing allogeneic hematopoietic stem cell transplantation (HSCT) ^2,3^, with genetic predisposition ^4^, or suffering from severe respiratory illnesses like Influenza or Covid-19 ^5^. IPA poses a major clinical challenge due to limited diagnostic tools, insufficient efficacy of available antifungal therapies, and increasing antifungal drug resistance ^3,6–8^. Hence, IPA is associated with high mortality rates ^2,9^ and poor long-term survival ^10^. Thus, understanding host defense and host pathogen-interaction are pivotal for the development of diagnostic biomarkers, targeted anti-virulence agents, and novel immunotherapies.

Several studies have shown that natural killer (NK) cells are an integral part of protective antifungal immunity ^11,12^. For instance, HSCT recipients with delayed NK-cell reconstitution or low NK-cell counts are at higher risk of developing IPA, indicating that NK cells are indispensable for fungal clearance ^13^. NK cells constitute 5-15% of the blood lymphocyte repertoire and can either secrete regulatory cytokines to recruit and stimulate other immune cells or elicit cytotoxicity directly against target cells ^14–16^. Therefore, after target recognition, the lytic immunological synapse between NK cells and the target cell is formed, enabling NK-cell-induced elimination of target cells through death receptor-mediated apoptosis ^17–20^. The central mechanism of target cell lysis is the delivery of cytotoxic molecules, such as perforin or granzyme B, which are stored in granules ^17–19,21^. Immunological synapse formation between NK cells and *A. fumigatus*, NK-cell degranulation, and the presence of LAMP1 (CD107a) surrounding perforin at the NK-cell surface following degranulation have been described in detail ^22^.

Innate immune cells, including NK cells, are commonly activated by stimulation of *pattern recognition receptors* (PRR). These receptors detect *pathogen-associated molecular patterns* (PAMPs) of invasive pathogens. Several fungal-reactive PRRs of NK cells have been described. For instance, the natural cytotoxicity receptor (NCR) NKp30 was shown to recognize β-1,3-glucan on the surface of *Candida albicans* and *Cryptococcus neoformans* ^23^, while the activating human NK-cell receptor NKp46 interacts with the *Candida glabrata* adhesins Epa1, Epa6, and Epa7 to trigger antifungal cytotoxicity ^24^. Furthermore, we previously identified the NK-cell receptor CD56 as a PRR recognizing *A. fumigatus*. Upon contact with *A. fumigatus* hyphae, CD56 relocalizes actin-dependently toward the NK-cell/hyphal interaction side, leading to prominent reduction of CD56-fluorescence intensity. Blocking of CD56 resulted in decreased *A. fumigatus*-induced NK-cell activation and chemokine secretion, suggesting a functional relevance of CD56 in this NK cell-hyphal recognition^25^. However, while PAMPs for the NK cell receptors NKp46 ^24^ and NKp30 ^23^ have been identified, the *A. fumigatus* ligand of CD56 has remained unknown.

PAMPs are commonly either microbial structural components (e.g., cell wall polysaccharides) or lipopolysaccharides and are often required for pathogen survival or virulence ^23,26,27^. The cell wall of *A. fumigatus* is a complex and dynamic structure, which alters in composition depending on the fungal morphotype and environmental cues ^28,29^. It predominantly comprises polysaccharides and, to a smaller extent, proteins, lipids, and pigments ^30,31^. The main polysaccharides in the cell wall of *A. fumigatus* hyphae include chitin, β-1,6-branched β-1,3-glucan, α-1,3-glucan, galactomannan, and galactosaminogalactan ^28,32–35^.

Given our prior observation that CD56 solely interacts with *A. fumigatus* germ tubes or hyphae but not with conidia, we specifically focused on hyphal PAMPs to identify the *A. fumigatus* ligand of CD56. Using a combination of purified cell-wall components, biochemical treatments, and *A. fumigatus* mutants, we herein identified galactosaminogalactan (GAG), especially de-*N*-acetylated GAG, as the fungal ligand of CD56. The GAG/CD56 interaction triggered strong NK-cell activation, along with potent release of cytotoxic effectors and immune-enhancing chemokines. Furthermore, supernatants of GAG-pulsed NK cells inhibited fungal growth and enhanced the anti-*Aspergillus* activity of PMNs, suggesting GAG as a potential immunotherapeutic target in IPA.

## Results

### CD56-mediated recognition of *A. fumigatus* depends on intact cell-wall polysaccharides but not proteins

In a first step, we sought to determine whether the binding partner of CD56 on *A. fumigatus* hyphae is a cell wall protein or polysaccharide. Therefore, we depleted proteins in the cell wall of *A. fumigatus* germ tubes by proteinase K treatment before co-culture with NK cells and flow cytometric quantification of CD56 expression after fungal stimulation. Relocalization of CD56 toward the immunological synapse with *A. fumigatus* is accompanied by a remarkable decrease in fluorescence intensity ^25^. While this effect was seen with untreated *A. fumigatus*, no increase in CD56 fluorescence intensity was found after proteinase K treatment (Figure 1A), suggesting that cell-wall-bound proteins are not the ligands of CD56.

**Figure 1.**
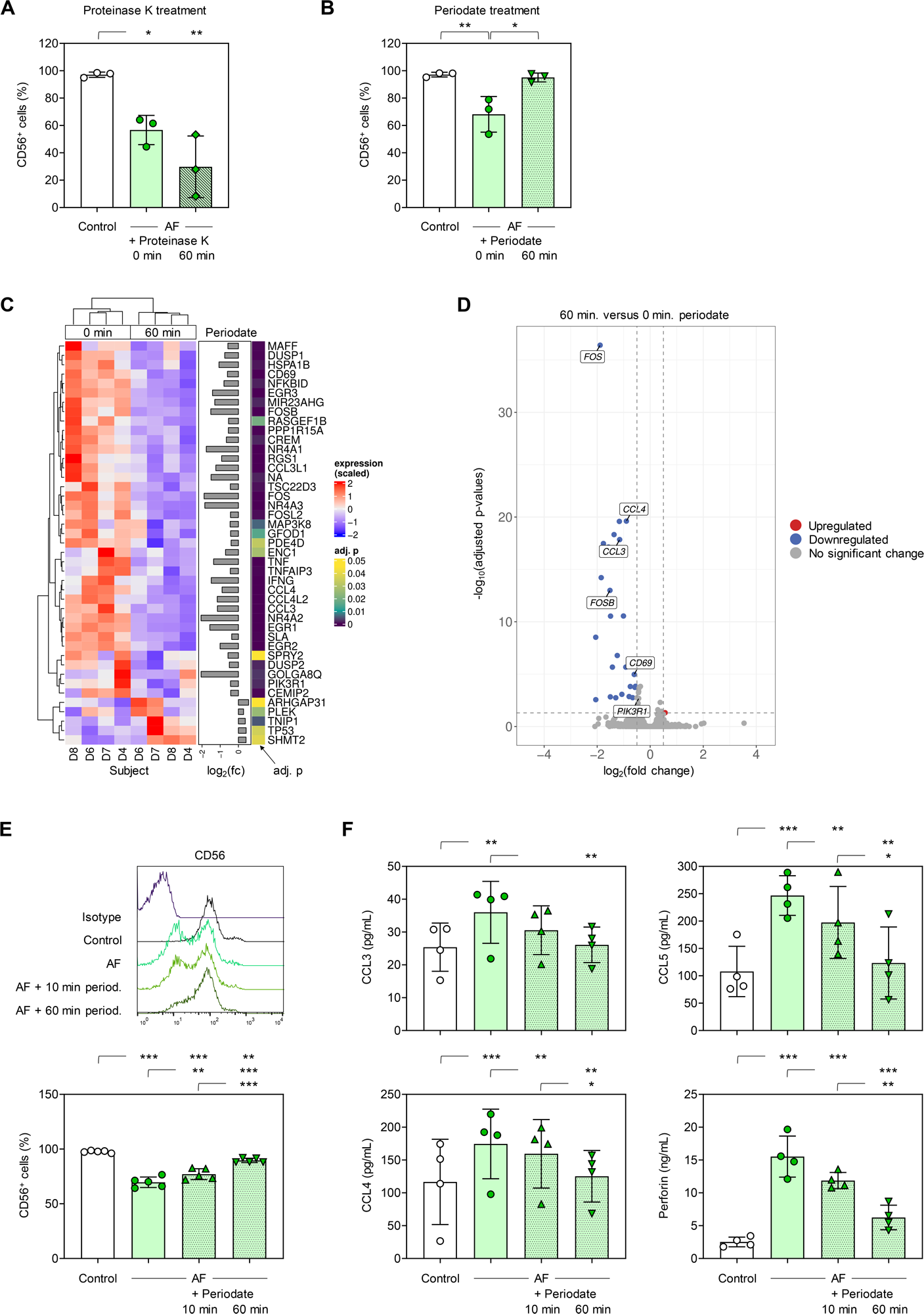
CD56 expression on naïve NK cells (Control) and NK cells stimulated for 3 h with *A. fumigatus* germ tubes, depending on proteinase K (A) or periodate (B) pre-treatment of the germ tubes. Data for 3 independent donors are shown. (C) Significantly differentially expressed genes after 3-h stimulation of NK cells with *A. fumigatus* germ tubes pre-treated (60 min) or not (0 min) with periodate. Abbreviations: adj. p = adjusted p-value, D = donor, fc = fold change. (D) Volcano plot summarizing transcriptional changes in NK cells stimulated with periodate-treated *A. fumigatus* germ tubes compared to cells stimulated with untreated germ tubes. Genes with a log2 fold change > 0.5 and an adjusted p-value < 0.05 are highlighted. Selected immune-related genes have been labelled. (E) CD56 expression on naïve NK cells (Control) and NK cells stimulated for 3 h with *A. fumigatus* germ tubes, depending on the duration of periodate pre-treatment of the germ tubes. Representative histograms and data for 5 independent donors are shown. (F) Chemokine and perforin release by naïve NK cells (Control) and NK cells stimulated for 3 h with *A. fumigatus* germ tubes, depending on the duration of periodate pre-treatment of the germ tubes. (A-B; E-F) Columns and error bars indicate means and standard deviations, respectively. Repeated measures one-way analysis of variance with Tukey’s post-hoc test. * p < 0.05, ** p < 0.01, *** p < 0.001.

Next, we treated *A. fumigatus* germ tubes with periodate that oxidizes cell wall polysaccharides having vicinal diols and co-cultured them with NK cells. Compared to NK cells co-cultured with untreated *A. fumigatus,* flow cytometry revealed no decrease in NK-cellular CD56-fluorescence intensity when challenged with periodate-treated germ tubes (Figure 1B). This suggest impaired CD56 relocalization of NK cells co-cultured with periodate-treated germ tubes. To reinforce these findings, bulk RNA sequencing was performed to identify genes that are differentially regulated in NK cells upon exposure to periodate-treated versus untreated *A. fumigatus* germ tubes. NK cells challenged with periodate-treated *A. fumigatus* showed 43 significantly differentially expressed genes compared to cells exposed to untreated germ tubes (Figure 1C). Out of these 43 genes, 38 genes (28 with a >0.5 log_2_ fold change) showed weaker expression in NK cells stimulated with periodate-treated fungus, including genes associated with NK-cell activation (e.g., *CD69*) and chemokine secretion (e.g., *CCL3* and *CCL4*) (Figure 1C-D).

Next, we confirmed that declined CD56 interaction of *A. fumigatus* after periodate treatment is dependent on the duration of periodate exposure (Figure 1E). Furthermore, periodate treatment time-dependently decreased the capacity of *A. fumigatus* germ tubes to elicit secretion of macrophage inflammatory protein (MIP)-1α (CCL3), MIP-1β (CCL4) and perforin from NK cells compared to untreated germ tubes (Figure 1F). Altogether, these results suggest that CD56 on human NK cells interacts with a hyphal cell wall polysaccharide and not a protein.

### Binding of CD56 to *A. fumigatus* cell wall galactosaminogalactan triggers NK-cell activation

To identify the specific cell wall polysaccharide that binds to CD56, we used an enzyme-linked immunosorbent assay (ELISA), in which different cell-wall polysaccharides extracted from the *A. fumigatus* hyphal cell wall were coated on microtiter plates and then incubated with recombinant CD56. Although CD56 interacts only with germ tubes and hyphae of *A. fumigatus*, we also tested binding to conidia-specific cell surface components, RodA protein (RodAp) and melanin pigment. CD56 was found to bind with high affinity to galactosaminogalactan (GAG), especially to its urea-insoluble fraction (PGG) enriched in *N*-acetylgalactosamine (Figure 2A) ^36^. GAG is a heteropolysaccharide composed of α-1,4-linked monomers of galactose, *N*-acetylgalactosamine (GalNAc), and galactosamine (GalN), synthesized during germination of *A. fumigatus* ^37–39^.

**Figure 2.**
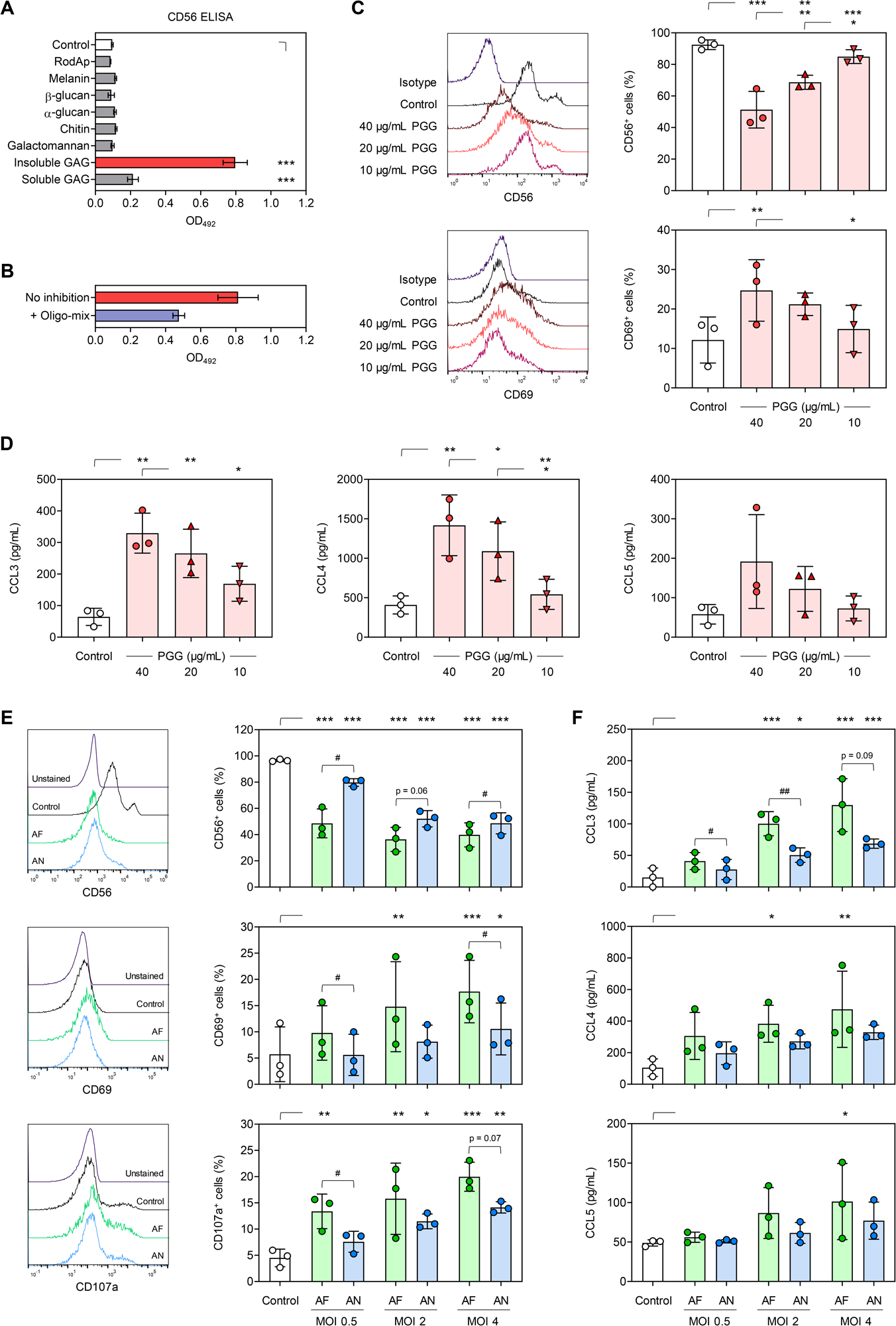
(A) CD56 binding to fungal carbohydrates and proteins, as determined by enzyme-linked immunosorbent assay. GAG = galactosaminogalactan, RodAp = surface rodlet protein/hydrophobin. N = 3 technical replicates. One-way analysis of variance (ANOVA) with Dunnett’s post-hoc test versus Control, i.e., no coating. (B) CD56 binding to GAG, with and without pre-incubation with GAG oligosaccharide fractions. N = 2 technical replicates. (C) CD56 and CD69 expression on naïve NK cells (Control) and NK cells stimulated for 24 h with different concentrations of urea-insoluble galactosaminogalactan (PGG). Representative histograms and data for 3 independent donors are shown. (D) Chemokine release by naïve NK cells (Control) and NK cells stimulated for 24 h with different concentrations of PGG. N = 3 independent donors. (C-D) Repeated measures (RM) one-way ANOVA with Tukey’s post-hoc test. (E) CD56, CD69, and CD107a expression on naïve NK cells (Control) and NK cells stimulated for 6 h with *A. fumigatus* (ATCC46645, AF) or *A. nidulans* (ATCC11267, AN) at different multiplicities of infection (MOIs). Representative histograms for one donor at MOI 4 are shown. (F) MOI-dependent stimulation of NK-cellular chemokine secretion after 6-h stimulation with AF or AN. (E-F) N = 3 independent donors. RM one-way ANOVA with Dunnett’s post-hoc test versus Control, i.e., unstimulated NK cells (asterisks). In addition, AF and AN stimulation at each MOI was compared using paired t-Test (hash signs). (A-F) Columns and error bars indicate means and standard deviations, respectively. */# p < 0.05, **/## p < 0.01, *** p < 0.001.

Next, we confirmed the specificity of the interaction between PGG and CD56. Therefore, PGG was coated on ELISA plates and probed with soluble CD56 that has been pre-incubated with acid-hydrolyzed PGG. Indeed, this hydrolysate markedly reduced the binding affinity of recombinant CD56 to PGG (Figure 2B), corroborating that the interaction between PGG and CD56 is specific.

To test whether PGG also interacts with CD56 on human NK cells, NK cells were stimulated with different concentrations of purified PGG. Using flow cytometry, we detected significant dose-dependent reduction in NK-cellular CD56 fluorescence after stimulation with purified PGG (Figure 2C). Additionally, PGG stimulation led to increased expression of the NK-cell activation marker CD69 (Figure 2C) and elicited strong secretion of cytotoxic effector molecules (granzyme B and perforin) and chemokines (CCL3, CCL4, and CCL5) from NK cells (Figures S1 and 2D).

Previous studies found significant differences in cell wall-bound GAG among *Aspergillus* species. Specifically, Lee *et al*. reported that *A. fumigatus* harbors the highest amount of cell wall-associated GAG, whereas *A. nidulans* produces low quantities of GAG ^39,40^. Therefore, we compared the decrease in CD56 expression on the NK cell surface and NK-cell activation after exposure to *A. fumigatus* and *A. nidulans* at different multiplicities of infection (MOIs). NK cells co-cultured with *A. fumigatus* showed stronger reduction in CD56 fluorescence intensity at all tested MOIs than NK cells confronted with *A. nidulans* (Figure 2E). Likewise, *A. fumigatus* induced stronger surface expression of CD69 and CD107a as well as chemokine secretion than *A. nidulans*, especially at a low MOI (Figure 2E and 2F). These results indicate that the quantity of cell wall-associated GAG plays a role in CD56-mediated NK-cell activation.

### CD56 binds to wild-type *A. fumigatus* hyphae but not to mutants lacking GAG

As purified GAG (i.e., PGG) from *A. fumigatus* triggered strong NK-cell activation and cytokine release, we next investigated the potential of the GAG-deficient mutant strains Δ*agd3* ^41^ and Δ*uge3* ^42^ to interact with CD56 on NK cells. Therefore, we co-incubated wildtype (WT) *A. fumigatus* (strain Af293) and the two otherwise isogenic mutants Δ*uge3* and Δ*agd3* (lacking key enzymes for GAG biosynthesis) with soluble CD56, followed by staining with fluorescent anti-CD56 antibody. WT *A. fumigatus* showed strong binding of CD56 to the hyphal surface, whereas CD56 did not bind to the hyphal surface of either GAG-deficient mutant (Figure 3A). To rule out any unspecific binding of the fluorescent antibody to hyphal structures, we co-incubated all strains with bovine serum albumin (BSA) protein and stained them with fluorescently-labeled anti-CD56 antibody. These control experiments showed only dim autofluorescence of the hyphae (Figure 3A right bottom image). In summary, these findings indicate specific binding of CD56 only to the cell wall of WT *A. fumigatus* hyphae that produce functional GAG.

**Figure 3.**
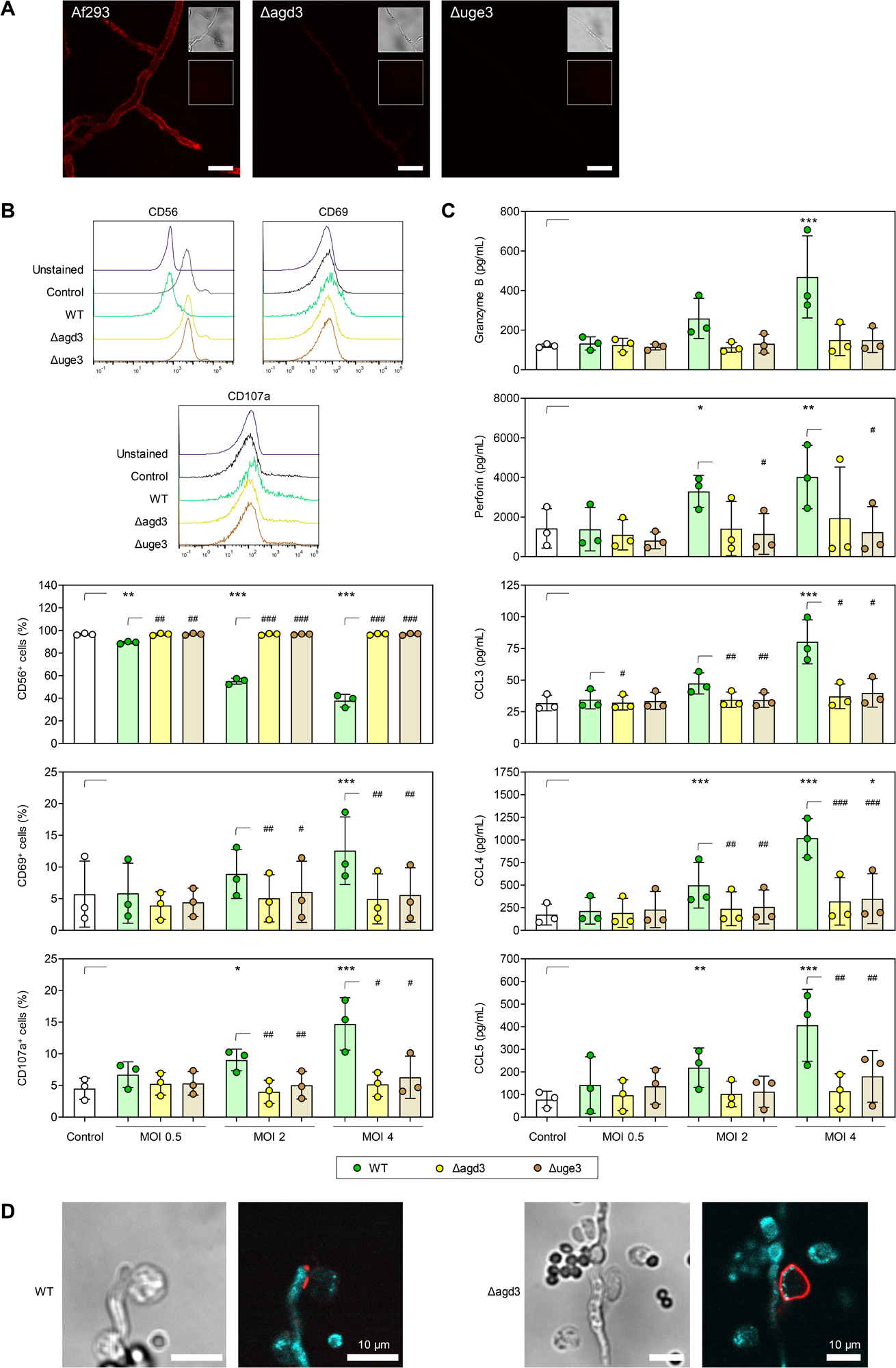
(A) Representative fluorescent micrographs (z-projection of 3-4 slices with 1 µm distance, representative dataset from ≥5 independent experiments) of hyphae of wild-type (WT) *A. fumigatus* Af293 and two galactosaminogalactan (GAG)-deficient *A. fumigatus* mutants (Δ*uge3* and Δ*agd3*) co-cultured with soluble CD56, followed by staining with fluorescent anti-CD56 antibody. The insets represent the bright-field image (top) corresponding to the confocal laser scanning microcopy (CLSM) image and a control image (bottom; BSA stained with anti-CD56 antibody) of the respective strain. Scale: 10 μm. (B) CD56, CD69, and CD107a expression on naïve NK cells (Control) and NK cells stimulated for 6 h with WT Af293 or the GAG-deficient *A. fumigatus* mutants (Δ*uge3* and Δ*agd3*) at different multiplicities of infection (MOIs). Representative histograms for one donor at MOI 4 are shown. (C) MOI-dependent induction of NK-cellular secretion of granzyme B, perforin, and chemokines after 6-h stimulation with WT, Δ*uge3*, and Δ*agd3* Af293. (B-C) N = 3 independent donors. Repeated measures (RM) one-way analysis of variance (ANOVA) with Dunnett’s post-hoc test versus Control, i.e., unstimulated NK cells (asterisks). In addition, results for stimulation with the 3 strains at each MOI was compared using RM one-way ANOVA with Dunnett’s post-hoc test versus WT (hash signs). Columns and error bars indicate means and standard deviations, respectively. */# p < 0.05, **/## p < 0.01, ***/### p < 0.001. (D) CLSM micrographs of NK cells co-cultured with WT Af293 and Δ*agd3 A. fumigatus* hyphae. CD56 was stained with anti-CD56 Alexa Fluor 647 (red) to assess the CD56 localization. Germ tubes could be detected via their auto-fluorescence (cyan). Scale: 10 μm.

### *A. fumigatus* mutants lacking GAG fail to elicit NK-cell activation

To further strengthen the role of GAG in CD56-mediated hyphal recognition and its potential to induce NK-cell activation, we co-cultured NK cells with WT *A. fumigatus* Af293 or the GAG-deficient mutants Δ*agd3* and Δ*uge3* at different MOIs. While NK cells confronted with the WT strain showed a marked and MOI-dependent reduction in CD56 fluorescence intensity, no decrease in CD56 expression was observed after co-culture with *A. fumigatus* strains lacking Agd3 and Uge3, even at an MOI of 4 (Figure 3B), indicating no accumulation of CD56 at the interaction side of NK cells with the mutant fungi.

Consistently, NK cells confronted with WT *A. fumigatus* hyphae but not with the Δ*agd3* or Δ*uge3* mutant strains showed an MOI-dependent increase in NK-cell activation (CD69) and degranulation (CD107a) (Figure 3B). Consequently, and in contrast to WT *A. fumigatus* hyphae, Δ*agd3* and Δ*uge3* mutant strains did not induce secretion of the cytotoxic effector molecules perforin and granzyme B as well as the chemokines CCL3, CCL4, and CCL5 (Figure 3C). This suggests that GAG is involved in *A. fumigatus*-induced, CD56-mediated NK-cell activation.

### GAG is indispensable for CD56 accumulation at the site of *A. fumigatus*/NK-cell interaction

To further corroborate that CD56 on NK cells binds to *A. fumigatus* GAG, we incubated NK cells with WT *A. fumigatus* hyphae or hyphae of the GAG-deficient strain Δ*agd3* and performed confocal laser scanning microscopy (CLSM) to assess the CD56 localization. Consistent with our previous study ^25^, NK cells co-cultured with the WT strain displayed a strong CD56 signal at the contact site, whereas NK cells exposed to the deacetylase-deficient Δ*agd3* mutant maintained a homogenously distributed CD56 fluorescence signal on the plasma membrane (Figure 3D) ^41^. This confirms that CD56 relocalizes to the fungal interface upon binding of its ligand GAG.

### Deacetylated residues of GAG are crucial for interaction with CD56 and NK-cell activation

Given the absence of CD56 binding to Δ*agd3* mutant hyphae, we further sought to test the importance of deacetylation on the CD56-GAG interplay. We first analyzed CD56 binding to either fully *N*-acetylated GAG (aPGG), fully de-*N*-acetylated GAG (dePGG), and native GAG (PGG), using ELISA. In contrast to native GAG, no binding of aPGG to CD56 was observed, whereas dePGG showed intermediate binding to CD56 (Figure 4A). Incubation of PGG, aPGG, and dePGG with only the secondary antibody or a combination of anti-CD56 and secondary antibody in the absence of recombinant CD56 yielded only a weak background signal, confirming that binding of CD56 to (de)PGG is specific (Figure 4A).

**Figure 4.**
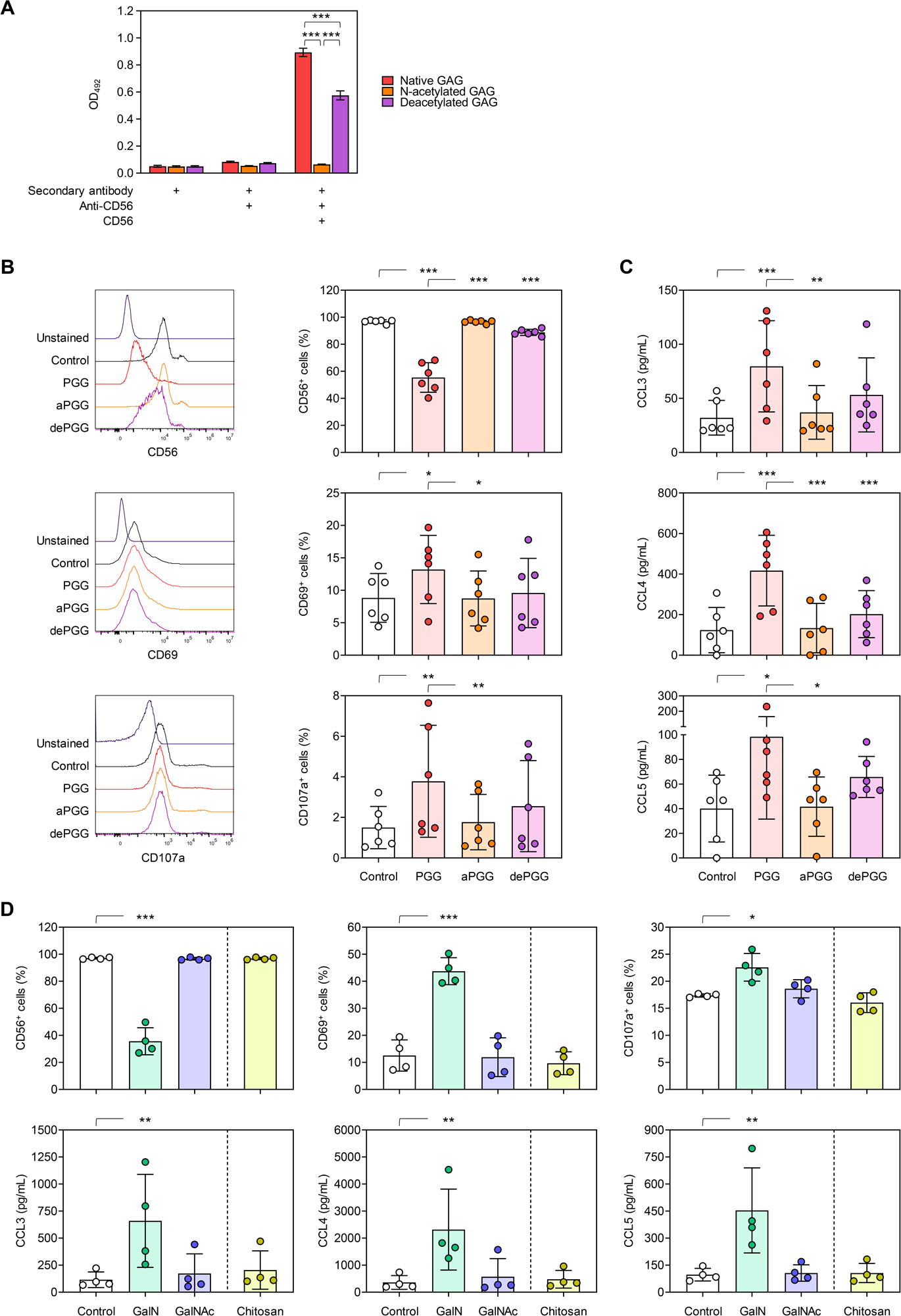
(A) CD56 binding to fully acetylated (aPGG), fully deacetylated (dePGG), and native galactosaminogalactan (PGG), as determined by enzyme-linked immunosorbent assay (ELISA). Additional conditions with incomplete ELISA setup were included to preclude unspecific binding of the secondary antibody or anti-CD56 to the carbohydrates. N = 3 technical replicates. One-way analysis of variance (ANOVA) with Tukey’s post-hoc test was performed for each assay setup. (B) CD56, CD69, and CD107a expression on naïve NK cells (Control) and NK cells stimulated for 24 h with PGG, aPGG, or dePGG. Representative histograms for one donor are shown. (C) NK-cellular chemokine secretion after 24-h stimulation with PGG, aPGG, or dePGG. (B-C) N = 6 independent donors. Repeated measures (RM) one-way ANOVA with Tukey’s post-hoc test. (D) CD56, CD69, and CD107a expression on naïve NK cells (Control) and NK cells stimulated for 24 h with GalN oligomers, GalNAc oligomers, or chitosan. NK-cellular chemokine secretion after 24-h stimulation with GalN oligomers, GalNAc oligomers, or chitosan. N = 4 independent donors. RM one-way ANOVA with Tukey’s post-hoc test. (A-D) * p < 0.05, ** p < 0.01, *** p < 0.001.

Next, we assessed the capacity of aPGG, dePGG, and native PGG to activate human NK cells. Consistent with the ELISA results, we observed a significant decrease in CD56-fluorescence intensity of NK cells stimulated with native PGG but not with aPGG, while dePGG elicited a modest decrease in CD56-fluorescence positivity. Similarly, stimulation of NK cells with native PGG caused a strong upregulation of CD69 and CD107a expression (Figure 4B), and triggered significant secretion of CCL3, CCL4, and CCL5 (Figure 4C). NK cells incubated with dePGG but not with aPGG showed modest degranulation and secreted slightly higher levels of CCL3, CCL4 and CCL5 than unstimulated NK cells (Figure 4C). Collectively, these findings indicate that the degree of deacetylation of GAG plays a role for full CD56-mediated NK cell activation.

To further elucidate the specificity of the interaction of CD56 with deacetylated galactosamine residues of GAG, we stimulated NK cells with GAG oligomers exclusively consisting of GalN or GalNAc residues. Also, considering the involvement of galactosamine in the interaction with CD56, we investigated the CD56 binding affinity of chitosan, a polymer composed of de-*N*-acetylated glucosamine and *N*-acetyl-glucosamine units ^43,44^. Both, GAG and chitosan are cationic polymers due to their deacetylation ^41,43,44^. Interestingly, only NK cells pulsed with GalN oligosaccharides displayed a significant reduction in CD56 fluorescence intensity, along with upregulation of CD69 and CD107a surface expression (Figure 4D). In contrast, GalNAc oligomers were unable to interact with CD56 and induce NK-cell activation. Likewise, chitosan showed neither binding to CD56 nor modulation of CD69 or CD107a (Figure 4D). Consequently, and in contrast to GalN oligomers, GalNAc and chitosan did not stimulate the secretion of chemokines CCL3, CCL4, and CCL5 (Figure 4D). Together, these findings indicate that the presence of galactose residues is not required for the interaction of CD56 with GAG. Moreover, the biochemically similar, positively charged polymer chitosan did not trigger NK-cell activation, further corroborating the specificity of the interaction between CD56 and GAG.

### The complete, GalN-rich GAG molecule is required for interaction of CD56 with *A. fumigatus*

To further validate that the complete, GalN-rich GAG molecule is required for CD56-mediated NK-cell activation, we preincubated germ tubes of WT *A. fumigatus* (Af293) or Δ*agd*3 with the GAG-specific carbohydrate-active enzymes Sph3 (hydrolase degrading GalNAc homo-oligomers ^45^), Ega3 (hydrolase degrading GalN homo-oligomers ^46^), and/or Agd3 (deacetylating homo-oligomers ^41,47^). As shown above, without enzyme pretreatment, only WT *A. fumigatus* resulted in a prominent reduction in NK-cellular CD56 fluorescence intensity (Figure 5A). Addition of either of the hydrolases or a combination of both to the NK-*A. fumigatus* co-culture alleviated (WT) *A. fumigatus*-induced CD56 relocalization, corroborating CD56 binding to the complete, GalN-rich GAG molecule. Expectedly, pretreatment of Δ*agd3* germ tubes with Agd3, allowing for external complementation of the mutation and GAG deacetylation, significantly decreased NK-cellular CD56 fluorescence intensity to a similar extent as non-pretreated WT *A. fumigatus*. As seen with WT *A. fumigatus*, this effect can be alleviated by addition of hydrolases. As previously observed, all conditions that led to decreased CD56 fluorescence intensity in turn induced enhanced CD69 expression (Figure 5A) and triggered strong production of the cytotoxic effector molecules perforin and granzyme B (Figure 5B). Altogether, these data suggest that the intact, GalN-rich GAG molecule is required for CD56-mediated NK-cell activation and release of antifungal effector molecules.

**Figure 5.**
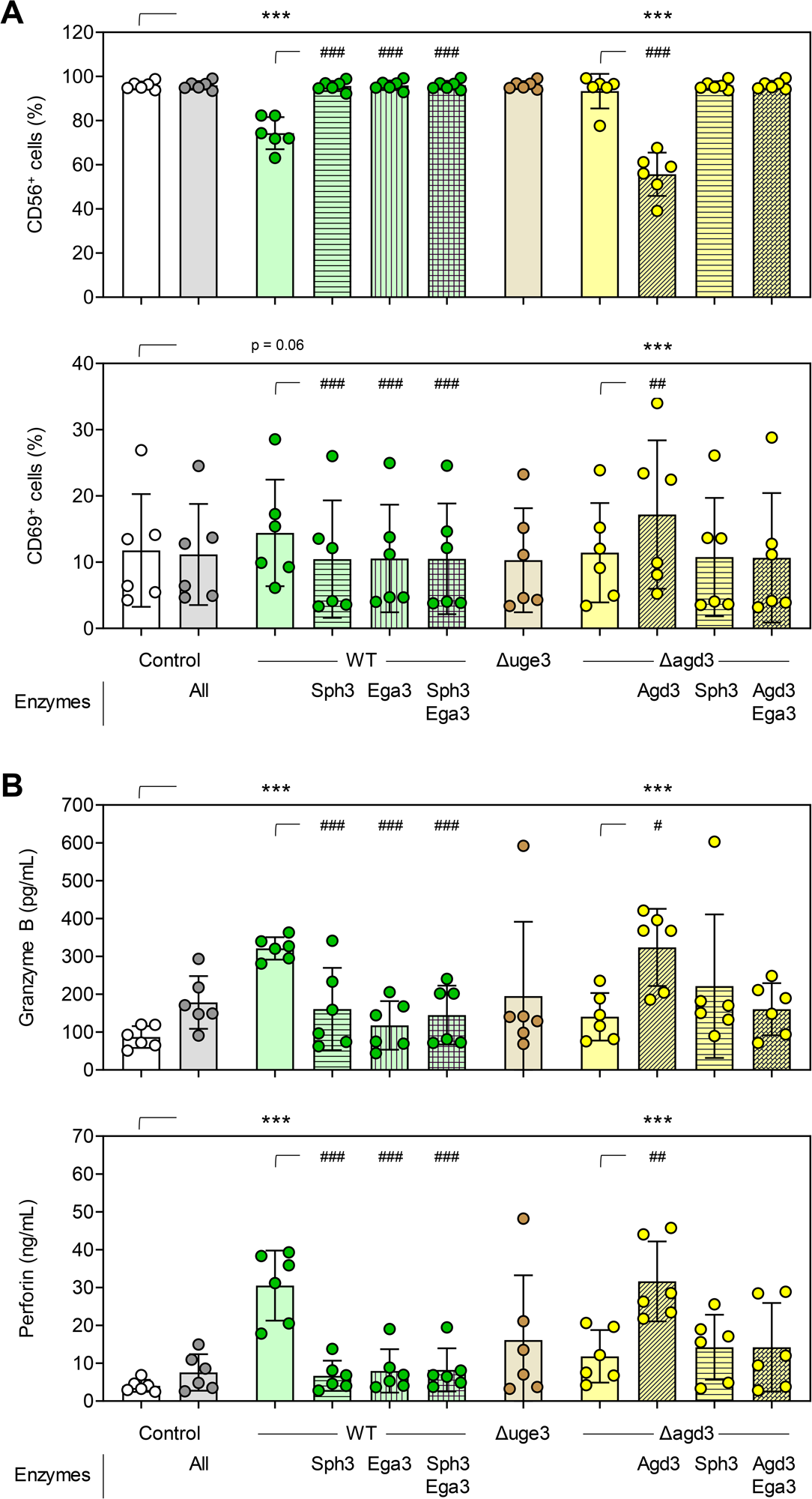
(A) CD56 and CD69 expression of unstimulated NK cells (Control) and *A. fumigatus*-stimulated (6 h) NK cells depending on the fungal strain (wild type [WT] Af293 or isogenic mutants with defective GAG biosynthesis) and its enzymatic pre-treatment. (B) Secretion of granzyme B and perforin by unstimulated NK cells (Control) and *A. fumigatus*-stimulated (6 h) NK cells depending on the fungal strain and its enzymatic pre-treatment. (A-B) N = 6 independent donors. Repeated measures (RM) one-way analysis of variance (ANOVA) with Dunnett’s post-hoc test versus Control, i.e., unstimulated NK cells (asterisks). Additionally, enzymatic pre-treatments of WT *A. fumigatus* and the Δ*agd3* mutant, respectively, were compared using RM one-way ANOVA with Dunnett’s post-hoc test versus no enzymatic pre-treatment (hash signs). Columns and error bars indicate means and standard deviations, respectively. */# p < 0.05, **/## p < 0.01, ***/### p < 0.001.

### Supernatants of PGG-stimulated NK cells inhibit fungal growth and stimulate polymorphonuclear neutrophils (PMNs)

Next, we sought to assess whether the effector responses induced by CD56/GAG interaction has any anti-*A. fumigatus* activity. Therefore, we used an IncuCyte time-lapse fluorescence microscope and the NeuroTrack image processing algorithm ^48^ to monitor fungal growth and hyphal branching. Compared to *A. fumigatus* alone, supernatants of NK cells stimulated with PGG significantly inhibited hyphal growth and branching (Figure 6A). In contrast, supernatants of unstimulated NK cells from the same donors did not inhibit fungal growth and morphogenesis (Figure 6A).

**Figure 6.**
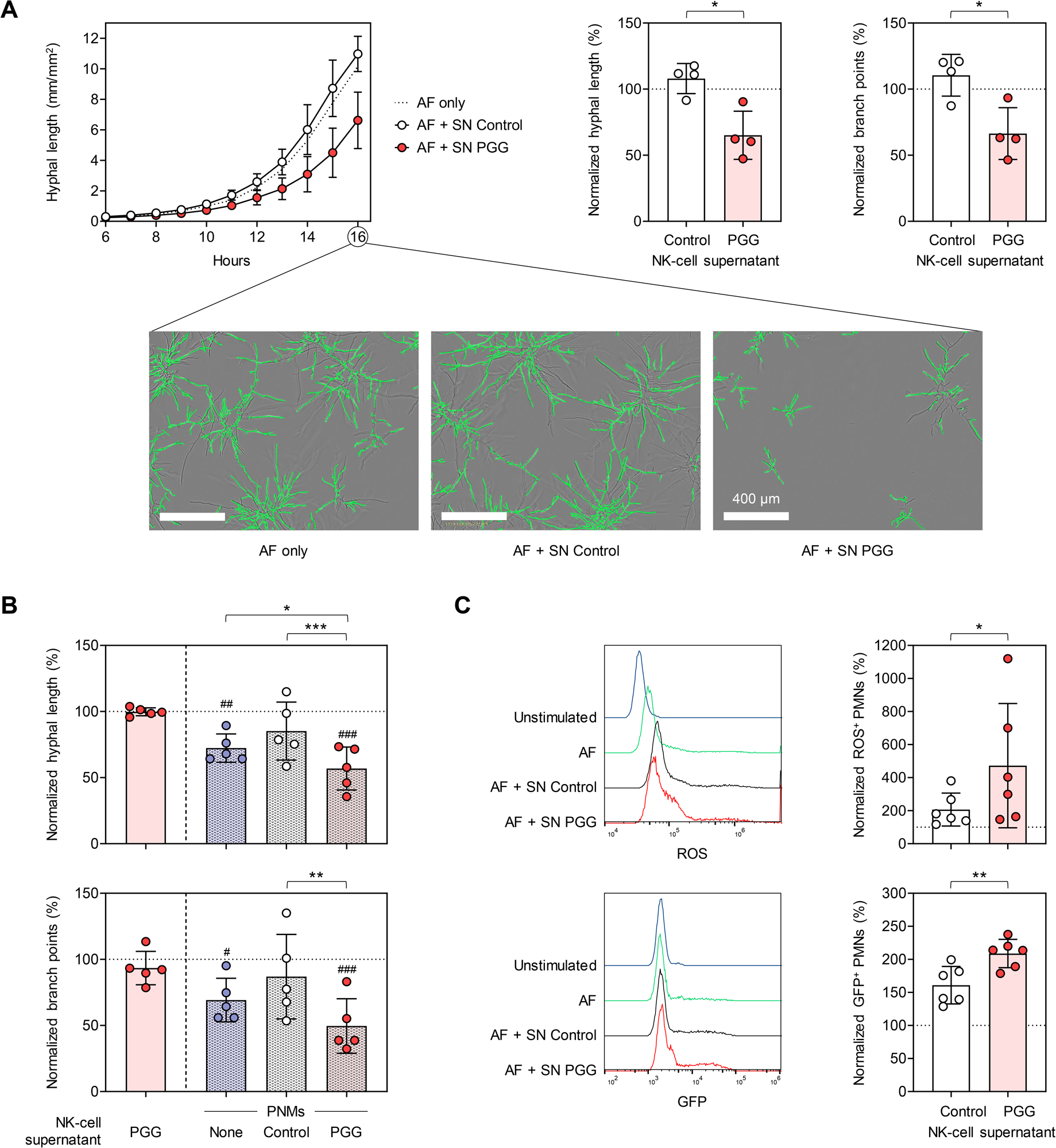
(A) Growth and morphogenesis of ATCC46645-GFP germlings in the presence of supernatants from unstimulated NK cells (SN Control) or PGG-stimulated NK cells (SN PGG) was quantified using the IncuCyte NeuroTrack assay and compared to ATCC46645-GFP grown without NK-cell supernatants (AF only). Left panel: Kinetics of hyphal length during transition from germlings to hyphae (6-16 h of culture). Center and right panel: Hyphal length and branch point numbers at 16 h of culture, normalized to the “AF only” control (dashed line, 100%). The assay was performed using NK-cell supernatants from n = 4 independent donors. Paired ratio t-test. In addition, images from a representative donor are shown. Green overlays indicate hyphal detection by the NeuroTrack algorithm. (B) Growth and morphogenesis of ATCC46645-GFP germlings in the presence of polymorphonuclear neutrophils (PMNs), with or without addition of SN Control or SN PGG, was quantified using the IncuCyte NeuroTrack assay. Hyphal length and branch point numbers at 16 h of culture were normalized to ATCC46645-GFP grown without PMNs and NK-cell supernatants (AF only, dashed line, 100%). AF grown without PMNs but with SN PGG was used as an additional control. N = 5 pairs of PMNs and NK-cell supernatants. Repeated measures (RM) 1-way analysis of variance and Dunnett’s post-hoc test versus “AF only” (hash signs) and “AF + PMNs without supernatant” (asterisks) were used to determine statistical significance of the combined effect of PMNs + supernatants and the impact of the supernatant on PMN-mediated fungal growth inhibition, respectively. (C) Production of reactive oxygen species (ROS) and phagocytosis was assessed upon co-culture of PMNs with *A. fumigatus* in the presence of SN Control or SN PGG for 3 h (ROS production) or 1 h (phagocytosis), respectively. Results were normalized to PMNs stimulated with *A. fumigatus* in the absence of NK-cell supernatant. N = 6 pairs of PMNs and NK-cell supernatants. Paired ratio t-test. (A-C) Columns and error bars indicate means and standard deviations, respectively. * p < 0.05, ** p < 0.01, *** p < 0.001.

In addition to their direct inhibitory effect on fungal growth, we evaluated the capacity of supernatants from PGG-activated NK cells, which contain increased levels of neutrophil-activating chemokines CCL3 and CCL4, to engage PMNs and enhance their anti-*Aspergillus* activity. Expectedly, PMNs partially inhibited fungal growth and hyphal branching of *A. fumigatus* in our IncuCyte NeuroTrack assay. Interestingly, a 2.5-fold lower concentration of supernatants from PGG-stimulated NK cells than in our previous experiment (Figure 6A), a concentration that *per se* did not inhibit mycelial proliferation, significantly enhanced suppression of hyphal growth and morphogenesis by PMNs (Figure 6B).

Additionally, we analyzed *A. fumigatus*-induced ROS production and phagocytosis of *A. fumigatus*-GFP conidia. While supernatants of unstimulated NK cells did not impact neutrophilic ROS response to the germ tubes, PGG-pulsed NK-cell supernatants significantly enhanced *Aspergillus*-induced ROS release of PMNs (Figure 6C). Consistently, supernatants of PGG-pulsed NK cells had a stronger stimulatory effect on phagocytosis of *A. fumigatus*-GFP by PMNs than supernatants of unstimulated NK cells (Figure 6C). Altogether, supernatants of PGG-stimulated NK cells elicited both direct antifungal activity and an indirect immunostimulatory effect through engagement of PMNs.

## Discussion

While alveolar macrophages and PMNs form the first line of antifungal defense in the lung, NK cells are an integral part of second-line defense against *A. fumigatus*. Several studies showed an involvement of CD56 in NK-cell-mediated cytotoxicity ^49–51^. For instance, the deletion of CD56 resulted in failed polarization during immunological synapse formation ^52^. We have previously shown that CD56 serves as a PRR for *A. fumigatus* that is required for proper antifungal activity of NK cells ^25^. Although the identity of fungal CD56 ligands remained unclear, it was known that only *A*. *fumigatus* hyphae but not resting conidia are capable to activate NK cells ^53^. Herein, we identified GAG as the PAMP of *A. fumigat*us hyphae that interacts with CD56 on NK cells, thereby activating NK cells and inducing potent secretion of chemokines and cytotoxic effector molecules, mirroring previous findings upon *A. fumigatus* stimulation ^25,53^. Notably, GAG-deficient mutants or *A. fumigatus* hyphae pre-treated with GAG-depleting hydrolases failed to elicit NK-cell activation and secretion of cytotoxic mediators and chemokines, underscoring the functional relevance of the CD56/GAG interaction.

GAG was reported to be located at the surface of the hyphal cell wall and profusely secreted in the extracellular matrix ^42^ where it plays a critical role in host-pathogen interactions ^37^. This polysaccharide is considered a multifunctional virulence factor contributing to biofilm formation ^42^, host cell adhesion ^41,42,54^, immune evasion ^28^, epithelial cell damage, and platelet activation. Moreover, the level of cell wall GAG production is directly associated with the varying virulence within the *Aspergillus* genus. ^39,42,55–57^. However, thus far, no mammalian receptor for GAG was known.

Purified native GAG bound to CD56 and elicited pronounced NK-cell activation, ruling out that impaired NK-cellular interaction with GAG-deficient hyphae was only due to their impaired adherence ^41,42,54^. Moreover, we were able to specifically visualize the interaction of CD56 with hyphal cell wall structures only in GAG-producing *A. fumigatus* but not in GAG-deficient mutants, further corroborating that GAG is essential for hyphal targeting by NK cells.

As Δ*agd3 A. fumigatus*, which produces GAG that lacks deacetylated GalN residues and therefore cannot stick to the hyphal surface ^41^, did not show binding to CD56, GAG composition appears to be crucial for its recognition by CD56. Given that both Δ*agd3 A. fumigatus* and purified fully *N*-acetylated PGG showed no interaction with CD56, we conclude that the presence of GalN in GAG is required for the binding of CD56 to GAG. Although both native PGG and fully de-*N*-acetylated PGG displayed binding to CD56, the interaction with fully de-*N*-acetylated PGG was significantly weaker, possibly due to steric hindrance. Purified GalN oligomers but not GalNAc oligomers displayed CD56-binding-affinity and elicited NK cell activation and chemokine secretion. This finding is consistent with prior reports that the degree of acetylation is an important modulator of the immunostimulatory versus anti-inflammatory properties of the exopolysaccharide GAG ^41,58^. Specifically, Briard *et al.* highlighted the role of deacetylated GAG for NLRP3 inflammasome activation through translational inhibition and induction of endoplasmic reticulum stress, in turn promoting protective immunity ^55^. Deacetylation of acetylated residues renders polymers cationic and introduces biologically unique properties, including adhesion to anionic surfaces such as the hyphal cell wall, plastic, and host cell membranes ^41^. However, even though both GAG and chitosan are de-*N*-acetylated polymers with cationic properties ^43,44^, chitosan did not exhibit immunostimulatory activity on NK cells, corroborating the specificity of the CD56/GAG interaction.

Recently, Picard *et al*. reported distinct effects of CD56 on primary NK-cell function and concluded that CD56 induces NK-cell degranulation, IFN-γ secretion, and morphological changes ^59^. Interestingly, that study did not reveal any major impact of CD56 on NK-cellular cytotoxicity. In our setting, we demonstrate that direct CD56/GAG interaction contributes to the release of antifungal effector molecules and cytotoxicity, resulting in the inhibition of hyphal growth, suggesting that GAG-activated NK cells may exert this effect by releasing perforin to reduce hyphal metabolic activity ^53^. Other NK-cellular effector functions that were not captured by our experiment with NK-cell supernatants, e.g., death receptor-mediated apoptosis, might also play a role. However, these effects are difficult to study, because the GAG-deficient mutants have altered growth and adhesion characteristics, making unbiased side-by-side comparisons in co-culture experiments difficult. Our data also showed that chemokine secretion from NK cells upon GAG stimulation leads to enhanced neutrophil activation and engagement of neutrophils into the antifungal immune response. While stimulating NK cells, GAG has been previously shown to contribute to pulmonary immunopathology by promotion of neutrophil apoptosis ^36,60^, induction of IL1RA secretion ^58,61^, and activation of the macrophage inflammasome ^55^. It is conceivable that NK-cellular cytotoxicity is crucial to overcome GAG-induced immunopathology. This hypothesis could explain the observation that poor NK-cell reconstitution significantly increased IPA risk in allogenic HSCT recipients, even in the absence of other cytopenia ^13^.

Thus far, detailed CD56 downstream signaling is largely unknown but is likely mediated by different activation kinases such as Syk, PI3K, and Erk ^59^. Therefore, functional studies in combination with multimodal CD56 downstream pathway analyses will be an important future direction. Unfortunately, no CD56 knockout mouse model is available, as no immediate CD56 homologue is present in murine NK cells. Therefore, future studies using CD56-deficient human NK cells, e.g., by means of CRISPR/Cas9 to specifically delete CD56, would be needed. Our study is subject to further limitations, including the partial lack of functional data due to the absence of an *in vivo* model for CD56 and the sole use of isolated NK cells without consideration of their complex interplay with other immune cell populations or tissue. Lastly, because this study focused on identifying and characterizing the fungal ligand of CD56, we did not include other fungal PAMPs and human PRRs in our analyses.

In summary, our data shed new light on the distinct *A. fumigatus*-NK cell interaction by providing inaugural evidence that GAG is an activating ligand of CD56 on human NK cells. To our knowledge, in this study, we have identified the first microbial ligand recognized by this receptor. Detailed insights into the specific immunomodulatory cellular signatures and downstream pathways involved in the CD56/GAG interaction could help to develop new alternative antifungal immunotherapies involving NK cells (e.g., chimeric antigen receptor NK cells). Moreover, further insights into the specific immunoregulatory properties and virulence attributes associated with specific biochemical patterns of the GAG molecule could enable the development of new targeted anti-virulence strategies and vaccines against *A. fumigatus*.

## Acknowledgements

The authors wish to thank all blood donors.

The study was supported by the Deutsche Forschungsgemeinschaft (DFG) within the Collaborative Research Center CRC124 FungiNet “Pathogenic fungi and their human host: Networks of interaction,” DFG project number 210879364 (project A2 to UT, HE and JL, C3 to OKu, and INF to SS and GP). We thank Jean-Paul Latgé, Institute of Molecular Biology and Biotechnology (IMBBFORTH), University of Crete, Heraklion, Greece, for his great support and advice in the preparation of this manuscript.

## Author Contributions

The study was conceived by LH, SW and JL. Experiments were planed and performed by LH, FN, NT, VA, SSWW, LS, EW and KH. Resources were provided by TF, FLM, DS, MV and PLH. Data were analyzed by LH, SS and SW. Data were visualized by SS and SW. Project administration and supervision were led by VA, TF, UT, FLM, DS, OKu, GP, SW, HE, and JL. Funding was acquired by HE, and JL. The original draft was written by LH, SW, and JL. All co-authors reviewed, edited, and approved the manuscript.

## Material and Methods

### Ethics statement

The processing of human peripheral venous blood from healthy adult donors was approved by the Ethics Committee of the University Hospital Würzburg (#302/12). Healthy volunteers donating venous whole blood for isolation of neutrophils gave informed written consent.

### Primary cell isolation

Peripheral blood mononuclear cells (PBMCs) were isolated from leukoreduction chambers obtained from plateletpheresis donations of healthy individuals. PBMCs were isolated by ficoll-histopaque^®^ (Sigma-Aldrich, St. Louis, MO, USA, 1.077 g/ml) density centrifugation. After separation, PBMCs were washed twice with 50 mL of HBSS buffer (Sigma Aldrich, St. Louis, MO, USA) supplemented with 2 mM EDTA (Sigma Aldrich, St. Louis, MO, USA) and 1% FCS (Sigma Aldrich, St. Louis, MO, USA). Thereafter, PBMCs were counted with a Vi-Cell XR counter (Beckman Coulter, Brea, USA). NK cells were isolated from PBMCs by negative selection using the human NK cell isolation kit (Miltenyi Biotec, Bergisch Gladbach, Germany). NK cells were cultured in Roswell Park Memorial Institute medium (RPMI, Gibco, Thermo Fisher Scientific, Waltham, MA, USA; 1×10^6^ cells/mL) supplemented with 10% heat-inactivated fetal calf serum (FCS, Sigma-Aldrich, St. Louis, MO, USA) and 120 µg/ml gentamicin (Refobacin, Merck, Darmstadt, Germany) at 37 °C and 5% CO_2_. Prior to co-culture experiments, NK cells were stimulated overnight with 1000 U/ml IL-2 (Proleukin® S, Novartis, Basel, Schweiz).

### Isolation of Polymorphonuclear Granulocytes (PMNs)

PMNs were isolated from 18 mL venous EDTA-anticoagulated blood from healthy donors using PolymorphPrep (ProteoGenix, Schiltigheim, France) gradient centrifugation at 590 ×*g* for 30 min. The interphase, containing PMNs, was carefully removed and washed with HBSS for 5 min at 590 ×*g*. To lyse remaining erythrocytes, 5 mL erythrocyte lysis buffer (EL buffer, Qiagen, Hilden, Germany) was added and PMNs were centrifuged, counted with a hemocytometer, and resuspended in RPMI + 10% FCS. Purity of the PMN suspension was analyzed by flow cytometry (CD3-PerCP^-^/CD66b-FITC^+^/CD14-PE^low^ cells > 90%, antibodies from Miltenyi Biotec, Bergisch Gladbach, Germany).

### Culture of fungal strains and preparation of germ tubes

*A. fumigatus* strains used in this study included Af293, Δ*agd3* mutant ^41^, Δ*uge3* mutant ^42^ (kindly provided by Donald Sheppard McGill University, Montreal, Quebec, Canada), as well as American Type Culture Collection (ATCC) reference strain 46645 and ATCC46645-GFP. In addition, *A*. *nidulans* strain ATCC11267 (Leibniz Institute, DSMZ-German Collection of Microorganisms and Cell Cultures, DSM820) was used. *A. fumigatus* and *A. nidulans* strains were plated on beer wort agar (Oxoid, Wesel, Germany) and incubated at 35 °C until conidiophores were visible. Conidia suspensions were prepared by rinsing the plates with sterile distilled water and passed through a 20-µm cell strainer (Miltenyi Biotec, Bergisch Gladbach, Germany) to eliminate residual mycelium. To generate germ tubes, 2×10^7^ conidia were incubated in RPMI medium (20 mL in 50 mL tubes) at 25 °C while shaking (200 rpm) until small germ tubes were visible. Germ tubes were centrifuged at 5,000 ×*g* for 10 min and resuspended in RPMI medium (4×10^6^/mL).

### Production of GAG-modifying enzymes

Glycoside hydrolases and deacetylase (gift from the Howell lab, Sick Kids, Toronto), were produced and purified as previously reported (Agd3 ^41,47^, Sph3 ^45,62^, Ega3 ^46^).

### Periodate/ Proteinase K treatment of *A. fumigatus* germ tubes

*A. fumigatus* ATCC 46645 conidia were seeded in 6-well plates (2×10^5^ conidia per well in 2 mL of RPMI medium) and incubated for 18 h at 30 °C. The supernatant was removed, and germ tubes were treated with 2 mL of 10 mM sodium *m*-periodate (Sigma Aldrich, St. Louis, MO, USA) in RPMI, or 50 µg/mL Proteinase K (Roche, Basel, Switzerland) for 60 min in RPMI for 10 min and 60 min, respectively, or were left untreated. Thereafter, wells were washed thrice by replacing two thirds of the supernatant with HBSS buffer supplemented with 2 mM EDTA and 1% FCS. After the last washing step, the washing buffer was completely removed, and 1 mL of pre-warmed RPMI containing 10% FCS was added to the hyphae. Three million NK cells were added to each well in a final volume of 3 mL per well. *A. fumigatus*/NK cell co-cultures were incubated for 3 h at 37 °C, 5% CO_2_ and subsequently analyzed by flow cytometry. Additionally, the effect of periodate treatment was analyzed by bulk RNA-Seq.

### CD56 binding to *A. fumigatus* cell wall polysaccharides

Polysaccharides of *A. fumigatus* mycelium were extracted and purified as previously reported. Briefly, galactomannan was isolated from mycelium membrane ^63^. GAG was purified from culture supernatant^58^. Chitin, β-1,3-glucans and α-1,3-glucans were extracted from cell wall after several chemical treatments ^64–66^. Polysaccharides were coated on 96-well microtiter plates (10 μg/well) overnight at room temperature (RT). Wells were blocked with PBS (Gibco, Thermo Fisher Scientific, Waltham, MA, USA) containing 1% BSA (Sigma Aldrich) at RT for 1 h, followed by incubation with recombinant CD56 protein (10 mg/mL,CliniSciences, Nanterre, France) in PBS + 1% BSA at RT for 1 h. Wells were washed thrice with PBS + 0.05% Tween 20 (Sigma Aldrich, St. Louis, MO, USA) and incubated with anti-CD56 antibody (1:50 in PBS + 1% BSA, CliniSciences, Nanterre, France) at RT for 1 h. After three washing steps with PBS + 0.05% Tween 20, wells were incubated with anti-mouse IgG peroxidase-conjugated (1:1000 in PBS + 1% BSA, Sigma Aldrich, St. Louis, MO, USA) at RT for 1 h, followed by an additional washing step. *O*-phenylenediamine dihydrochloride (Sigma Aldrich, St. Louis, MO, USA) was added as substrate. Color development was stopped with 4% H_2_SO_4_ (Sigma Aldrich) after 20 min. Optical density was measured at 492 nm as a surrogate of CD56 binding. Three independent experiments were performed.

### Inhibition of CD56 binding by GAG oligosaccharides

GAG was subjected to acid hydrolysis with 2 M HCl (Gibco, Thermo Fisher Scientific, Waltham, MA, USA), 100 °C for 3 h to obtain soluble oligosaccharides of various length (max. 25–28 monosaccharide units), as previously described ^36,58^. PGG (10 μg/well) was coated on a 96-well microtiter plate overnight at RT. Recombinant CD56 was pre-incubated with acid-hydrolyzed GAG oligosaccharides at RT for 30 min. The mixture was centrifuged, and the supernatant was added to the PGG-coated microtiter plate and incubated at RT for 1 h. Primary (anti-CD56 antibody 1:50 in PBS + 1% BSA, CliniSciences, Nanterre, France) and secondary (anti-mouse IgG peroxidase-conjugated, 1:1000 in PBS + 1% BSA, Sigma Aldrich, St. Louis, MO, USA) antibodies were added at RT for 1 h. *O*-phenylenediamine dihydrochloride (Sigma Aldrich, St. Louis, MO, USA) was used as substrate to measure absorbance at 492 nm, as described above. Two independent experiments were performed.

### NK-cell stimulation with purified GAG, its oligomers, or chitosan

To analyze binding of PGG to CD56 and PGG-induced NK-cell activation, IL-2-pre-stimulated NK cells (2×10^5^ cells/well) were incubated with different concentrations (40 µg/mL, 20 µg/mL, 10 µg/mL) of PGG ^36^ or with 10 µg/mL fully de-*N*-acetylated PGG or *N*-acetylated PGG ^58^ for 24 h at 37 °C and 5% CO_2_. For one experimental series, IL-2-pre-stimulated NK cells (2×10^5^ cells/well) were pulsed with 40 µg/mL GalN or, GalNAc oligomers ^58^ or chitosan (Sigma-Aldrich St. Louis, MO, USA) for 24 h at 37 °C and 5% CO_2_. For flow cytometric analysis of degranulation, brefeldin A (4 µg/mL, Sigma-Aldrich St. Louis, MO, USA), BD GolgiStop™ (0.67 µL/mL, BD Biosciences, San Jose, CA, USA), and CD107a-PE (0.5 µL/mL, Miltenyi Biotec, Bergisch Gladbach, Germany) were added after 6 h of incubation. A combination of 25 ng/mL phorbol-12-myristate-13-acetate (PMA, Sigma-Aldrich St. Louis, MO, USA) and 1 µg/mL ionomycin (Sigma-Aldrich St. Louis, MO, USA) served as positive control. NK-cell activation was analyzed by flow cytometry and ELISA, as described below.

### NK-cell infection assays

IL-2-pre-stimulated NK cells (2×10^5^ cells) were incubated with *A*. *fumigatus* (WT, Δ*agd3*, or Δ*uge3*) or *A. nidulans* germ tubes at MOIs of 0.5, 2, or 4 for 6 h at 37 °C, unless indicated otherwise. For one experimental series (see Figure 5), *A. fumigatus* germ tubes were pre-incubated with either 2 µM Sph3 hydrolase ^45^, 1 µM Ega3 hydrolase ^46^, 0.1 µM Agd3 deacetylase ^47^, or a combination of these enzymes for 30 min, followed by co-culture with NK cells at an MOI of 0.5. All tested co-culture conditions are summarized in Supplementary Table S1. For flow cytometric analysis of degranulation, brefeldin A (4 µg/mL, Sigma-Aldrich St. Louis, MO, USA), BD GolgiStop™ (0.67 µL/mL, BD Biosciences, San Jose, CA, USA), and CD107a-PE (0.5 µL/mL, Miltenyi Biotec, Bergisch Gladbach, Germany) were added after 1 h of incubation.

### Stimulation of PMN oxidative burst and phagocytosis by NK-cell supernatants

To generate culture supernatants, 2×10^5^ NK cells were incubated with 40 µg/mL PGG for 24 h in a 200-µL volume. NK cells were centrifuged at 300 ×*g* for 10 min; supernatant was harvested and stored at – 80 °C until further use. To assess oxidative burst, 2×10^5^ isolated PMNs (2×10^6^ cells/well) were incubated with *A. fumigatus* ATTC 46645 germ tubes (MOI 0.5) in a total of 25 µL RPMI + 10% FCS or seeded in an equivalent volume of RPMI + 10% FCS without fungal cells in a 96-well plate. Next, culture supernatant of NK cell-PGG co-cultures or NK cells cultured alone (40 µL, containing secreted metabolites of 40.000 NK cells) were added. The volume of each well was adjusted to 200 µL with cell-free RPMI + 10% FCS. A combination of 5 ng/mL PMA (Sigma-Aldrich St. Louis, MO, USA) and 0.2 µg/mL ionomycin (Sigma-Aldrich St. Louis, MO, USA) served as positive control. ROS production was analyzed by flow cytometry, as described below. To assess phagocytosis, 2×10^5^ PMNs were seeded in 96-well plate. Culture supernatant of NK cell-PGG co-cultures or NK cells cultured alone (40 µL, containing secreted metabolites of 40.000 NK cells) and *A*. *fumigatus* (ATCC46645-GFP) conidia were added at an MOI of 3 for 1 h at 37 °C. The volume of each well was adjusted to 200 µL with cell-free RPMI + 10% FCS. Phagocytosis rate was determined using flow cytometry.

### RNA isolation and bulk transcriptomic profiling

Total RNA from purified NK cells was isolated using QIAshredder columns and the RNeasy Mini Kit (Qiagen, Hilden, Germany) following the manufacturer’s protocol. RNA purity and concentration were tested with a NanoDrop ND-1000 spectral photometer (Thermo Fisher Scientific, Waltham, MA, USA). The integrity of RNA was determined with a 2100 Bioanalyzer (Agilent Technologies, Waldbronn, Germany) using RNA 6000 Pico or Nano LabChip Kits (Agilent Technologies, Waldbronn, Germany) according to the manufacturer’s instructions. RIN values of our samples were in the range of 7.3–10.0, indicating good quality of the RNA samples.

Library preparation was performed with the Illumina TruSeq® Stranded mRNA technology, according to the manufacturer’s protocol. RNA sequencing was performed by IMGM Laboratories GmbH (Martinsried, Germany) on the Illumina NextSeq® 500 next-generation sequencing system with 1×75 bp single-read chemistry. Raw files are accessible under the Gene Expression Omnibus accession number GSE241020.

### RNA-seq data processing and analysis

Preprocessing of raw reads, including quality control and gene abundance estimation, was performed with the GEO2RNAseq pipeline (v0.100.1) ^67^ in R version 3.5.1. Quality analysis was performed before and after trimming with FastQC (v0.11.7). Read-quality trimming was performed with Trimmomatic (v0.36). Adapter sequences were removed, window size trimming was performed (15 nucleotides, average Q < 25), 5′ and 3′ clipping was performed for any base with Q < 3, and sequences shorter than 30 nucleotides were removed. Reads were mapped against the human reference genome (GRCH 38, v109). First, the reference genome was indexed with exon information using HiSat2 (v2.1.0). Then, read alignment was performed using HiSat2 on the exon-indexed reference genome. Gene abundance estimation was performed with featureCounts (R package Rsubread, v1.34.0) in single-end mode with default parameters. MultiQC (v1.7) was used to summarize the output of FastQC, Trimmomatic, HiSat, featureCounts, and SAMtools. Count matrices were normalized using median-by-ratio normalization (MRN) as described before ^68^. Differential gene expression was analyzed by applying DESeq2 (v1.38.3) and adding donor origin as co-factor to the statistical design. Gene expression differences were considered significant at an adjusted p-value of ≤0.05. Hierarchical clustering was performed with the ward.D2 clustering method using MRN gene-abundance data with the R package ComplexHeatmap (v2.14.0).

### Flow cytometry

NK-cell activation and degranulation as well as neutrophil activation were assessed by flow cytometry. Therefore, NK-cell/PMN cultures were centrifuged for 10 min at 300 ×*g* at RT and cell pellets were resuspended in 100 µL HBSS supplemented with 2 mM EDTA and 1% FCS. Cells were stained extracellularly with fluorescent antibodies listed in Supplementary Table S2 and the fixable Viobility^TM^ Live/Dead Dye (Miltenyi Biotec, Bergisch Gladbach, Germany). After incubation in the dark for 15 min at 4 °C, cells were washed with 2 mL HBSS supplemented with 2 mM EDTA and 1% FCS, resuspended in 150 µL of 4% paraformaldehyde (Sigma-Aldrich, St. Louis, MO, USA), and incubated for 30 min at RT. Data were acquired on a FACSCalibur (Treestar/Becton & Dickinson, BD), Cytoflex flow cytometer (Beckman Coulter, Brea, CA, USA) or MACS Quant 10 flow cytometer (Miltenyi Biotec, Bergisch Gladbach, Germany). Downstream data analysis was performed with FlowJo10 (Treestar/Becton & Dickinson, BD) or Kaluza v.2.1 (Beckman Coulter, Brea, CA, USA). Gating strategies and representative raw data are shown in the Supplement.

### ELISA and multiplex cytokine assay

To quantify cytokine and chemokine release of stimulated NK cells, cell cultures were harvested and centrifuged at 300 ×*g* for 10 min. Culture supernatants were cryopreserved at -80 °C until further analysis. Concentrations of CCL3 (MIP-1α, R&D Systems, Minneapolis, MN, USA), CCL4 (MIP-1β, R&D Systems, Minneapolis, MN, USA), CCL5 (RANTES, Biolegend, San Diego, CA, USA), granzyme B (R&D Systems, Minneapolis, MN, USA), and perforin (abcam, Cambridge, UK) were determined using ELISA kits according to the manufacturer’s manual with minor modifications. Briefly, ELISAs were performed in 96-well half-area, high-binding plates with one-fourth of the volume recommended by the manufacturer. Absorbance was measured with a NanoQuant Infinite 200M Pro microplate reader (Tecan, Maennedorf, Switzerland). For multiplexed quantification of cytokine and chemokine concentrations in supernatants of NK cells infected with enzymatically pre-treated *A. fumigatus*, a ProcartaPlex™ 11-PLEX assay kit (ThermoFisher Scientific, Waltham, MA, USA) including GM-CSF, granzyme B, IFN-ɣ, IL-1α, IL-6, IL-8, CCL3, CCL4, perforin, CCL5, TNF-α was used according to the manufacturer’s instructions. Acquisition/measurement was performed using a Luminex detection system (Bio-Plex 200 system) and Bio-Plex Manager™ Software 6.2 (Bio-Rad, Hercules, CA, USA).

### Binding of soluble CD56 to *A. fumigatus* hyphae

Cellvis 8-well (Mountain View, CA, USA) coverslips were pre-coated with laminin (20 µg/mL, Sigma Aldrich, St. Louis, MO, USA) for 1 h at 33 °C and washed twice with ddH_2_O. *A. fumigatus* Af293, *Δagd3*, and *Δuge3* conidia were suspended in colorless RPMI (Gibco, Thermo Fisher Scientific, Waltham, MA, USA), added to the coated coverslips (5×10^4^ per well), and incubated at 33 °C until germination. Hyphae were incubated with soluble CD56 (5 µg/mL, R&D Systems, Minneapolis, MN, USA) or BSA (control, 5 µg/mL, Roth, Karlsruhe, Germany) in colorless RPMI (Gibco, Thermo Fisher Scientific, Waltham, MA, USA) for 2 h at 37 °C, 5% CO_2_. Wells were washed twice with colorless RPMI, fixed with 1% formaldehyde (Thermo Fisher Scientific, Waltham, MA, USA for 10 min, and washed once with HBSS (Sigma Aldrich, St. Louis, MO, USA). After blocking with 5% BSA (Roth, Karlsruhe, Germany) in HBSS (Sigma Aldrich, St. Louis, MO, USA) for 30 min, samples were stained with Alexa Fluor 647-labeled mouse-anti-human CD56 (10 μg/mL, Biolegend, San Diego, CA, USA) for 1 h and washed with HBSS (Sigma Aldrich, St. Louis, MO, USA). Stained samples were imaged with an LSM700 laser scanning confocal microscope (Carl Zeiss) with a plan-apochromat 63x/1.40 oil immersion objective. The following acquisition settings were used: image size, 71.35×71.35 μm; laser power, 8%; gain, 650; pinhole, 1 AU (53.7). Z-stacks were obtained at an interval of 1-2 µm. ImageJ/Fiji software was used for image processing.

### Fluorescence microscopy of NK-cell/*A. fumigatus* co-cultures

*A. fumigatus* Af293, *Δagd3*, and *Δuge3* (5×10^4^ conidia per well in colorless RPMI) were seeded on laminin-coated (20 µg/mL, Sigma Aldrich, St. Louis, MO, USA) 8-well coverslips and incubated overnight at 33 °C to enable germination, as described above. NK cells were added to the germ tubes at an MOI of 0.5 (1×10^5^ cells per well) and incubated for 4 h at 37 °C, 5% CO_2_. After incubation, samples were fixed with 1% formaldehyde (Thermo Fisher Scientific, Waltham, MA, USA) for 7 min, washed once with PBS (Gibco, Thermo Fisher Scientific, Waltham, MA, USA), and blocked with 2.5% BSA (Roth, Karlsruhe, Germany) in PBS (Gibco, Thermo Fisher Scientific, Waltham, MA, USA) for 30 min. Samples were stained immediately after fixation with Alexa Fluor 647-labeled mouse-anti-human CD56 (10 μg/mL, Biolegend, San Diego, CA, USA) for 1 h and washed with PBS (Gibco, Thermo Fisher Scientific, Waltham, MA, USA). CLSM images were recorded with a LSM700 system (Carl Zeiss) using a plan-apochromat 63 ×/1.40 oil immersion objective.

### IncuCyte imaging

To generate culture supernatants for IncuCyte experiments, 2×10^5^ NK cells were incubated with 40 µg/mL PGG for 24 h in a 200-µL volume. NK cells were centrifuged at 300 ×*g* for 10 min; supernatant was harvested and stored at – 80 °C until further use. For IncuCyte experiments, *A. fumigatus* spores (ATCC-GFP) were diluted in RPMI + 10% FCS to a final concentration of 3×10^3^/mL. Fifty microliters (150 conidia/well) were dispensed per well of a 96-well flat-bottom plate. To test the direct antifungal effect of PGG-pulsed NK cells, culture supernatant of NK cell-PGG co-cultures or NK cells cultured alone (37.5 µL, containing secreted metabolites of 37,500 NK cells) was added to the wells. The volume of each well was adjusted to 150 µL with cell-free RPMI + 10% FCS. To evaluate the capacity of PGG-stimulated NK cells to enhance fungal growth inhibition by PMNs, 1.5×10^4^ PMNs (effector/target ratio, 100) in 85 µL RPMI + 10% FCS were added to the wells. In addition, culture supernatant of NK cell-PGG co-cultures or NK cell cultured alone (15 µL, containing secreted metabolites of 15,000 NK cells) was added to the wells. The volume of each well was adjusted to 150 µL with cell-free RPMI + 10% FCS. In addition to biological replicates (n=5), each condition was tested in technical duplicates. IncuCyte microscopy and NeuroTrack-based image analysis (IncuCyte Zoom NeuroTrack software module) was performed according to Wurster *et al.* ^48^. In brief, well plates were imaged hourly in the IncuCyte Zoom HD/2CLR time-lapse microscopy system (Sartorius, Göttingen, Germany) equipped with an IncuCyte Zoom 10× Plan Fluor objective (Sartorius, Göttingen, Germany) for a period of 16 h. Acquisition time for the green channel was 400 ms. The following parameters were used for NeuroTrack analysis: neurite coarse sensitivity, 5; neurite fine sensitivity, 0.25; neurite width, 4 µm. Neurite length [mm/mm^2^] and numbers of branch points [1/mm^2^]) were compared to an “*A. fumigatus* only” control without NK-cell supernatants and/or PMNs.

