## Supplementary Figure, Supplementary Tables for "CD56-mediated activation of human natural killer cells is triggered by *Aspergillus fumigatus* galactosaminogalactan"

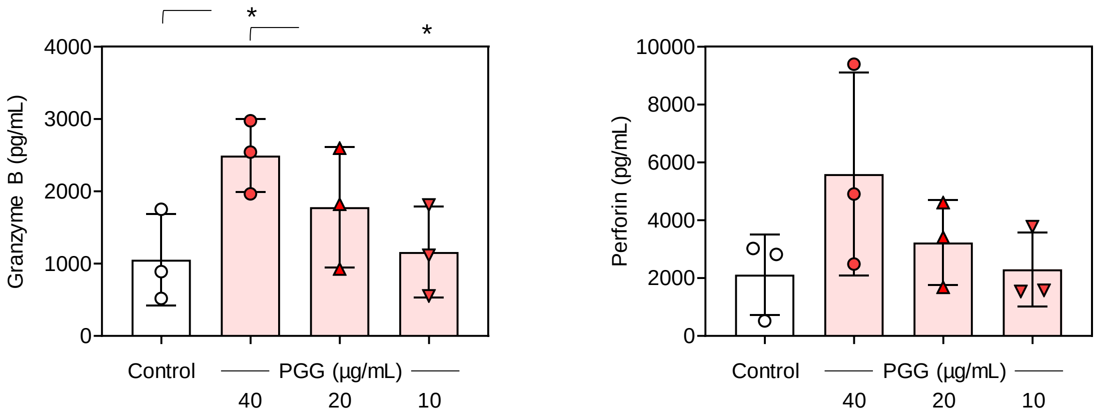


Figure S1

Release of granzyme B and perforin by naïve NK cells (Control) and NK cells stimulated for 24 h with different concentrations of urea-insoluble galactosaminogalactan (PGG). N = 3 independent donors. Columns and error bars indicate means and standard deviations, respectively. Repeated measures one-way ANOVA with Tukey’s post-hoc test. * $p < 0.05$.

Gating strategies

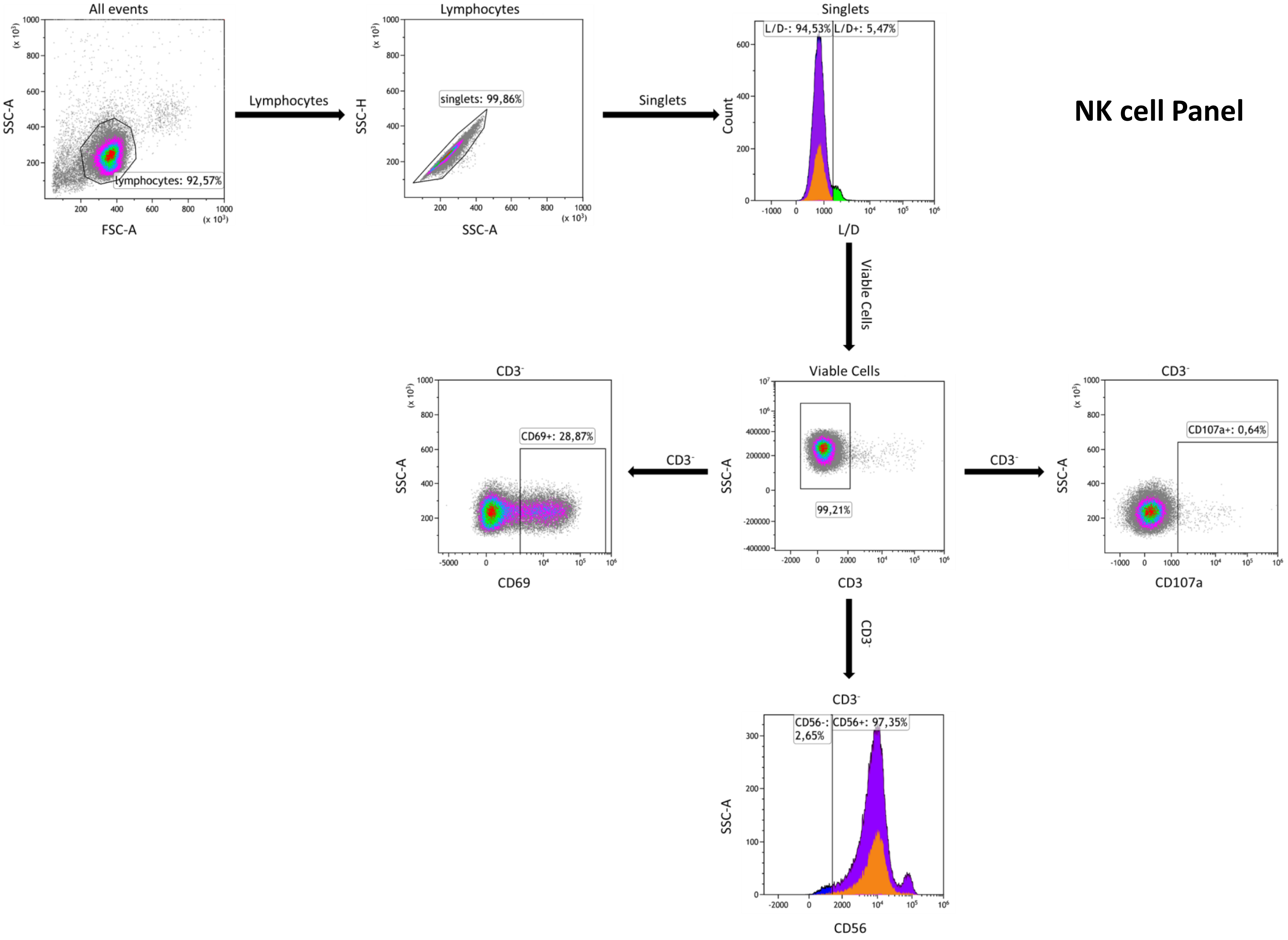

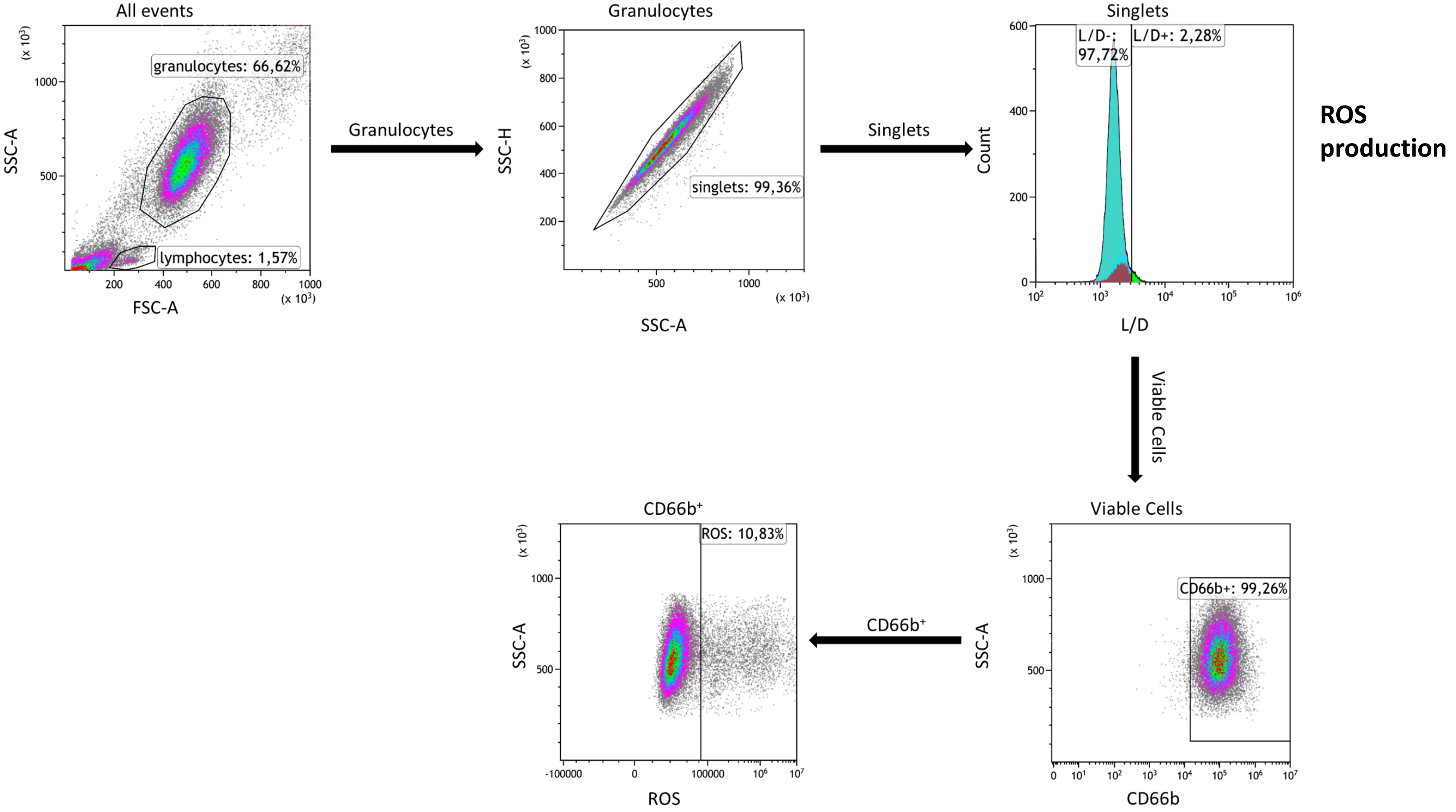
All events
granulocytes: 66,62%
lymphocytes: 1,57%
SSC-A
FSC-A
Granulocytes
Granulocytes
singlets: 99,36%
SSC-H
SSC-A
Singlets
Singlets
L/D-: 97,72%
L/D+: 2,28%
Count
L/D
ROS production
Viable Cells
Viable Cells
CD66b+: 99,26%
SSC-A
CD66b
CD66b⁺
CD66b⁺
ROS: 10,83%
SSC-A
ROS

### Phagocytosis

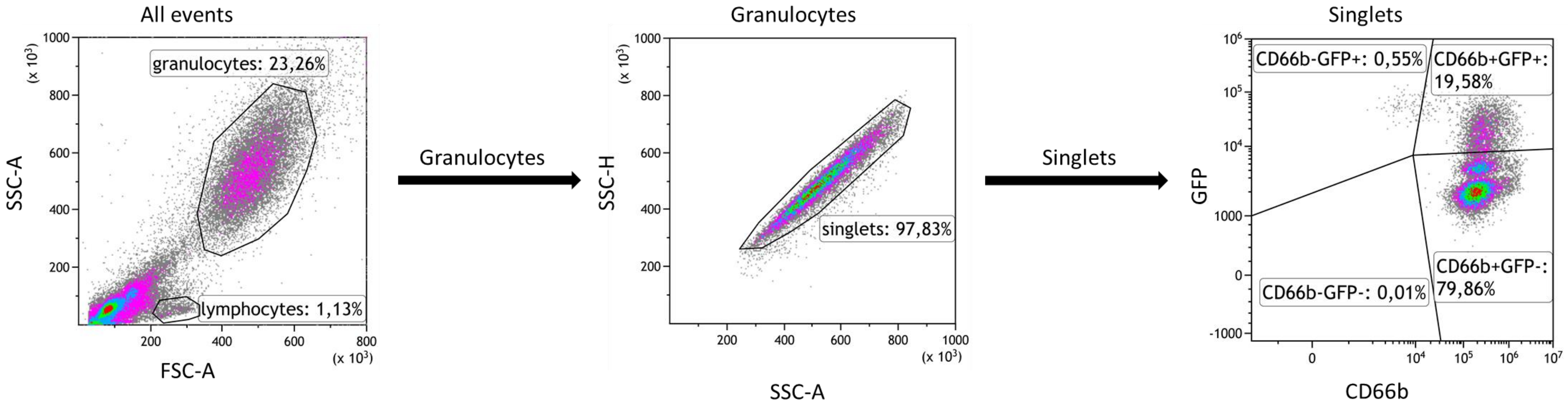


Gating strategy to study NK cell interaction, activation, and degranulation (1st figure)

Isolated Lymphocytes are selected according to their forward and side scattering and single lymphocytes were gated based on SSC-A, and SSC-H properties. Dead lymphocytes were excluded by Live/Dead staining. Natural killer (NK) cells were identified as $CD3^-$. Positive cells for NK cell surface marker CD56, for activation markers CD69, and degranulation marker CD107a were gated within each $CD3^-$ NK cell population. Fluorescence positivity was determined for all markers.

Gating strategy to determine ROS production by PMNs (2nd figure)

Debris was excluded from isolated granulocytes by light scatter properties and dead granulocytes were excluded by Live/Dead staining. Granulocytes were identified by CD66 expression and analyzed for ROS expression. Fluorescence positivity was determined for ROS.

Gating strategy to study phagocytosis (3rd figure)

Isolated granulocytes s are selected according to their forward and side scattering and single lymphocytes were gated based on SSC-A, and SSC-H properties. Conidia phagocytosed by PMNs are identified as $CD66b^+GFP^+$.

Supplementary Tables

Supplementary Table 1. Preparation of NK cell *A. fumigatus*-infection assay with wild type Af293 or Δ*agd3/* Δ*uge3* mutants and their enzymatic pre-treatment

| | NK cells | Af293 gt | Δ*uge3* gt | Δ*agd3* gt | Agd3 | Ega3 | Sph3 | PMA | Ionomycin | RPMI+10 % FCS |
|---|---|---|---|---|---|---|---|---|---|---|
| **Stock concentration** | $2*10^6$ cells/mL | $2*10^7$ gt | $2*10^7$ gt | $2*10^7$ gt | 132 µM | 596 µM | 568 µM | 1 mg/mL | 1 mg/mL | n/a |
| **Final concentration** | $1*10^6$ cells/mL | $4*10^6$ gt/mL | $4*10^6$ gt/mL | $4*10^6$ gt/mL | 0.1 µM | 1 µM | 2 µM | 25 ng/mL (pre-dilution 1:1000) | 1 µg/mL (pre-dilution 1:10) | n/a |
| **Unstimulated control (NK)** | 100 µL | | | | | | | | | 100 µL |
| **NK+Δ*uge3*** | 100 µL | | 25 µL | | | | | | | 75 µL |
| **NK+Af293** | 100 µL | 25 µL | | | | | | | | 75 µL |
| **NK+Af293+Sph3** | 100 µL | 25 µL | | | | | 25 µL | | | 50 µL |
| **NK+Af293+Ega3** | 100 µL | 25 µL | | | | 25 µL | | | | 50 µL |
| **NK+Af293+Ega3+Sph3** | 100 µL | 25 µL | | | | 25 µL | 25 µL | | | 25 µL |
| **NK+Δ*agd3*** | 100 µL | | | 25 µL | | | | | | 75 µL |
| **NK+Δ*agd3*+Agd3** | 100 µL | | | 25 µL | 25 µL | | | | | 50 µL |
| **NK+Δ*agd3*+Sph3** | 100 µL | | | 25 µL | | | 25 µL | | | 50 µL |
| **NK+Δ*agd3*+Agd3+Ega3** | 100 µL | | | 25 µL | 25 µL | 25 µL | | | | 25 µL |
| **NK+Agd3+Ega3+Sph3 (all enzymes)** | 100 µL | | | | 25 µL | 25 µL | 25 µL | | | 25 µL |
| **NK+PMA+Ionomycin** | 100 µL | | | | | | | 5 µL | 2 µL | 75 µL |

| | **NK cells** | **Af293 gt** | **Δ*uge3* gt** | **Δ*agd3* gt** | **Agd3** | **Ega3** | **Sph3** | **PMA** | **Ionomycin** | **RPMI+10 % FCS** |
|---|---|---|---|---|---|---|---|---|---|---|
| **Stock concentration** | 2*$10^{6}$ cells/mL | 2*$10^{7}$ gt | 2*$10^{7}$ gt | 2*$10^{7}$ gt | 132 µM | 596 µM | 568 µM | 1 mg/mL | 1 mg/mL | n/a |
| **Final concentration** | 1*$10^{6}$ cells/mL | 4*$10^{6}$ gt/mL | 4*$10^{6}$ gt/mL | 4*$10^{6}$ gt/mL | 0.1 µM | 1 µM | 2 µM | 25 ng/mL | 1 µg/mL | n/a |
| **Unstimulated control (NK cells)** | X | | | | | | | | | X |
| **NK+Δ*uge3*** | X | | X | | | | | | | X |
| **NK+Af293** | X | X | | | | | | | | X |
| **NK+Af293+Sph3** | X | X | | | | | X | | | X |
| **NK+Af293+Ega3** | X | X | | | | X | | | | X |
| **NK+Af293+Ega3+Sph3** | X | X | | | | X | X | | | X |
| **NK+Δ*agd3*** | X | | | X | | | | | | X |
| **NK+Δ*agd3*+Agd3** | X | | | X | X | | | | | X |
| **NK+Δ*agd3*+Sph3** | X | | | X | | | X | | | X |
| **NK+Δ*agd3*+Agd3+Ega3** | X | | | X | X | X | | | | X |
| **NK+Agd3+Ega3+Sph3 (all enzymes)** | X | | | | X | X | X | | | X |
| **NK+PMA+Ionomycin** | X | | | | | | | X | X | X |

X indicates that the compound was used for the respective condition. A total volume of 200 µL was used for all conditions.

Abbreviations: gt = germ tubes, PMA = Phorbol-12-myristate-13-acetate, RPMI = Roswell Park Memorial Institute medium, n/a = not applicable

Supplementary Table 2. Antibodies used for flow cytometric analyses

| **Name** | **Fluorochrome** | **Clone** | **Provider** | **Catalogue number** |
|---|---|---|---|---|
| CD3 | PE | UCHT1 | BD Pharmingen | 555333 |
| CD3 | PerCP | BW264/56 | Miltenyi Biotec | 130-113-131 |
| CD3 | APC-Vio770 | REA613 | Miltenyi Biotec | 130-113-136 |
| CD14 | APC-Vio770 | TÜK4 | Miltenyi Biotec | 130-113-144 |
| CD56 | FITC | REA196 | Miltenyi Biotec | 130-114-549 |
| CD56 | FITC | B159 | BD Pharmingen | 562794 |
| CD66b | PE-Vio770 | REA306 | Miltenyi Biotec | 130-119-768 |
| CD69 | PE-Vio615 | REA824 | Miltenyi Biotec | 130-112-617 |
| CD69 | PerCP | FN50 | Biolegend | 310928 |
| CD69 | PE-Vio770 | REA824 | Miltenyi Biotec | 130-112-615 |
| CD107a (LAMP-1) | PE | REA792 | Miltenyi Biotec | 130-111-621 |

All antibodies/dyes were used at 1% v/v, except for CD69 PercP (2% v/v), and CD3 PE (2% v/v)
